# Cytoplasmic flows caused by actomyosin contraction drive interkinetic nuclear migration

**DOI:** 10.1101/2020.04.09.034660

**Authors:** Amarendra Badugu, Andres Käch

**Author notes:** Correspondence to Amarendra Badugu.

## Abstract

Interkinetic nuclear migration (IKNM) is the process by which the nucleus migrates between apical and medial surfaces of pseudostratified epithelia. Previous studies have proposed force generating mechanisms, acting primarily on the nucleus. Having observed in drosophila wing discs that cytoplasmic components (lipid droplets and mitochondria) migrate alongside the nucleus, we used live imaging and particle tracking to demonstrate that the cytoplasm flows are responsible for the nucleus migration. We identify that nuclear migration in mitotic cells is preceded by a fast basal-to-apical flow of cytoplasm occurring over short time scales. We further show that, for the migration of basally located nuclei to an apical position, a slower flow of cytoplasm is responsible over a longer time scale. Our findings indicate that these flows are driven by acto-myosin contractile forces. These flows increase the hydrostatic pressure under the nucleus to exert a lifting force, much like a piston in a hydraulic cylinder.

## Introduction

Pseudostratified epithelial tissues (PSE) are monolayers of epithelial cells, arranged in a way that results in a multilayered appearance. PSE occur widely throughout the animal kingdom and form organ precursors (Norden, 2017). Examples include vertebrate neurogenic placodes which are regions of the embryonic ectoderm that generate the majority of cranial sensory systems (Graham et al., 2007), *danio rerio* embryonic retina (Norden et al., 2009), imaginal discs in *drosophila* (Meyer et al., 2011) and ectoderm in *nematostella* (Meyer et al., 2011). They are also found in retinal, cerebral and neural tube organoids before they differentiate (Norden, 2017). Every PSE cell extends from the apical surface to the basal lamina, with mitosis occuring at the apical side (Sauer, 1935). In the *drosophila* wing imaginal disc, the nucleus spends the interphase of the cell cycle in a region known as medial zone. Through a mechanism known as Interkinetic nuclear migration (IKNM), the nucleus migrates for mitosis into the mitotic zone (Fig. 1b-1b’, Fig. 1c), a 10 µm region at the apical surface (Meyer et al., 2011). If IKNM is perturbed, mitosis occurs basally (Meyer et al., 2011; Strzyz et al., 2015), leading to break-down of tissue order.

**Fig 1:**
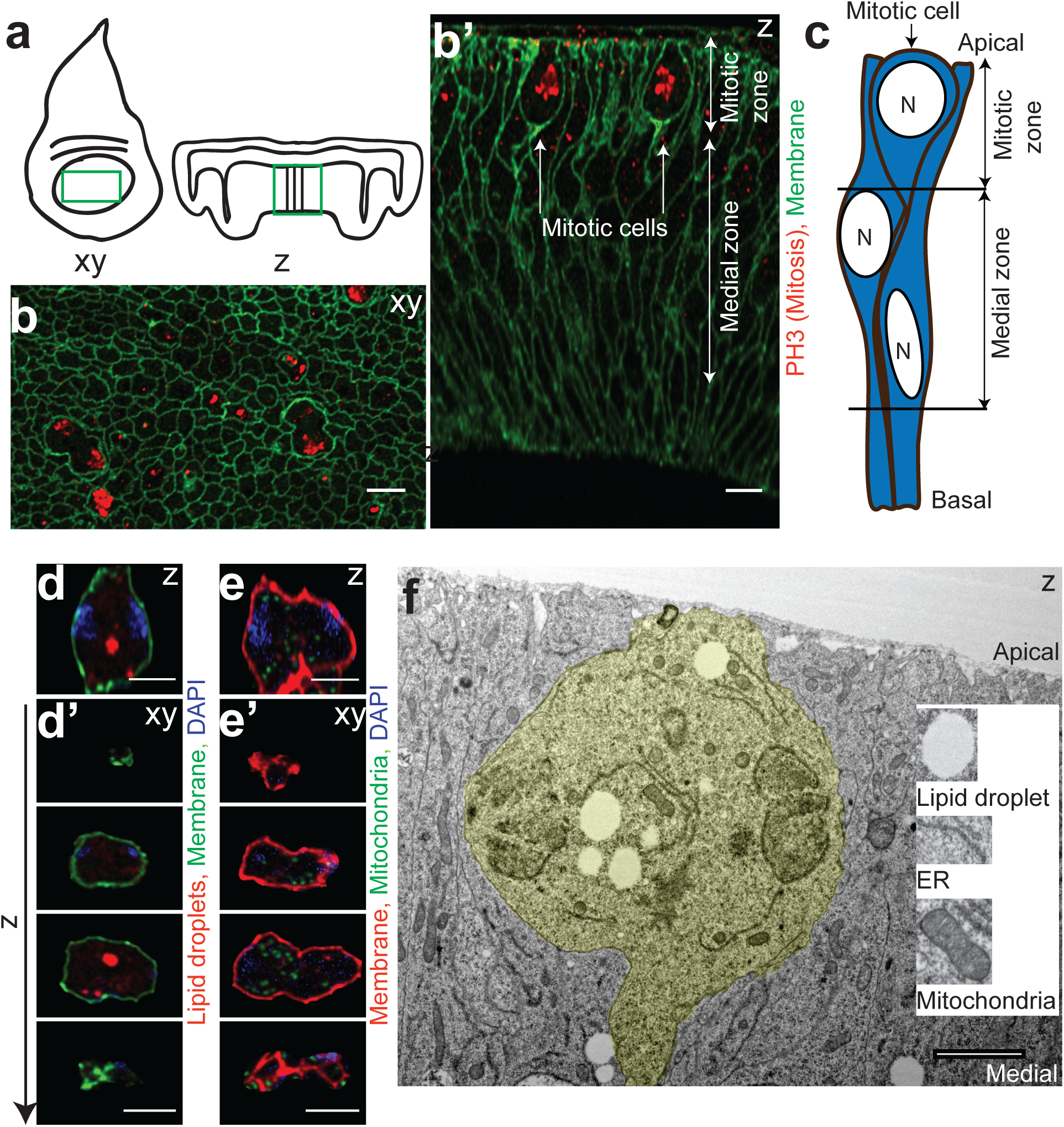
Mitotic cells are located at the apical surface in a PSE and contain cytoplasmic components. (a) Region of analysis in the wing imaginal disc. xy slice represents top-down view. z slice represents basolateral view. (b, b’) *c765-GAL4>UAS-PLCdelta-PH-GFP* with PH3 staining. (b) xy slice showing cells in different stages of mitosis. Prominent features are mitotic rounding and increased cell area. (b’) z slice showing mitotic zone where nucleus is present for cells undergoing mitosis and medial zone where nuclei are present in other phase of cell cycle. (c) Model representing the mitotic zone and medial zone.(d, d’) *c765-GAL4>UAS-PLCdelta-PH-GFP*, LipidTOX Deep Red and DAPI (d) z slice through middle of anaphase cell showing lipid droplets. (d’) xy slices of (d) 4um apart in z. (e, e’) *ap-GAL4* GAL80^ts^; UAS*-mito-HA-GFP>*UAS*-PLCdelta-PH-mcherry* and DAPI. (e) z slice through middle of late anaphase cell showing mitochondria (e’) xy slices of (e), 4um apart in z. (f) TEM image of a mitotic cell in anaphase with cytoplasmic components, labeled mitotic cell (yellow), lipid droplets, mitochondria and endoplasmic reticulum as in insert.. Scale bars: b-b’, d-d’, e’-e’: 5 µm, f: 2µm.

In the *danio rerio* embryonic retina, *drosophila* wing imaginal disc and in *nematostella* ectoderm, IKNM has been linked to t*h*e actin/myosin based cytoskeleton (Meyer et al., 2011; Norden et al., 2009). The exact mechanism by which the moving force is generated by actin/myosin is still not very fully understood (Norden, 2017) and further only the nuclear migration has been studied while the movement of cytoplasmic components has been largely neglected.

The cytoskeleton is a meshwork of filaments, motor proteins embedded in the cytoplasm (Nazockdast et al., 2017). The interactions between cytoplasm and the cytoskeleton (hydrodynamic interactions) have often been ignored though they can be quite useful in understanding force transduction mechanisms (Nazockdast et al., 2017). Since the cytoskeleton elements were embedded in a highly viscous cytoplasm, their interactions can instantaneously induce flows on the scale of the cell (Shelley, 2016).

Currently, cytoplasmic flows have been observed in a wide range of eukaryotic organisms such as plants, amoebae, nematodes (*C. elegans*) and *drosophila* often in unusually large cells such as the egg (Goldstein and van de Meent, 2015). It has been postulated that cytoplasmic flows have been evolved as effective mechanism in overcoming the slowness of diffusion in transporting organelles and metabolites over long length scales (100 µm-10 cm) (Goldstein and van de Meent, 2015). In *C. elegans*, long distance intracellular flows generated by cortical tension were responsible for polarization of the zygote (Mayer et al., 2010). In *drosophila* embryos, cytoplasmic flows from the apical constriction of the tissue drives cell elongation (He et al., 2014). The motion of the nucleus during IKNM has been postulated as being an indirect consequence of a cytoplasmic flow generated by contractions of an actomyosin ring though this remains to be shown (Lee and Norden, 2013). By tracking cytoplasmic components that could reveal the cytoplasmic flows, we investigated whether such a mechanism exists in the wing imaginal disc.

## Results

Cytoplasmic components can be used as a proxy for studying the displacement of cytoplasm, i.e cytoplasmic flows. We examined the distribution of organelles in the mitotic cells and non-dividing cells in the wing pouch. We observed from confocal data that lipid droplets and mitochondria were present in cells undergoing mitosis (Fig. 1d-d’, 1e-e’, Supplementary Movie. S1). In the non-dividing cells, lipid droplets were most abundant in basal regions (Supplementary Fig. S1a-b’’). Mitochondria on the other hand were present along the entire apical-basal axis of cells (Supplementary Fig. S1b-b’’). We prepared and analyzed electron micrographs of wing imaginal discs and confirmed that mitotic cells contained lipid droplets and mitochondria (Fig. 1f). Lipid droplets were significantly larger in size and fewer in number than mitochondria making them a good marker for particle tracking. Additionally, groups of tightly packed lipid droplets moving together as lipid droplet clusters (LDC) provide larger structures that can be tracked easily.

### Type I apical directed flow precedes mitosis

We performed live imaging of the pouch region in the wing imaginal disc (Fig. 1a). We examined mitotic events and tracked the motion of LDC along the apical-basal axis back in time. The tracking data revealed the existence of a cytoplasmic flow, starting at the basal surface and directing the LDC towards the apical surface (Fig. 2a, Supplementary Fig. S2c, Supplementary Movie. S2). We define this apical-directed flow as type I flow.

**Fig 2:**
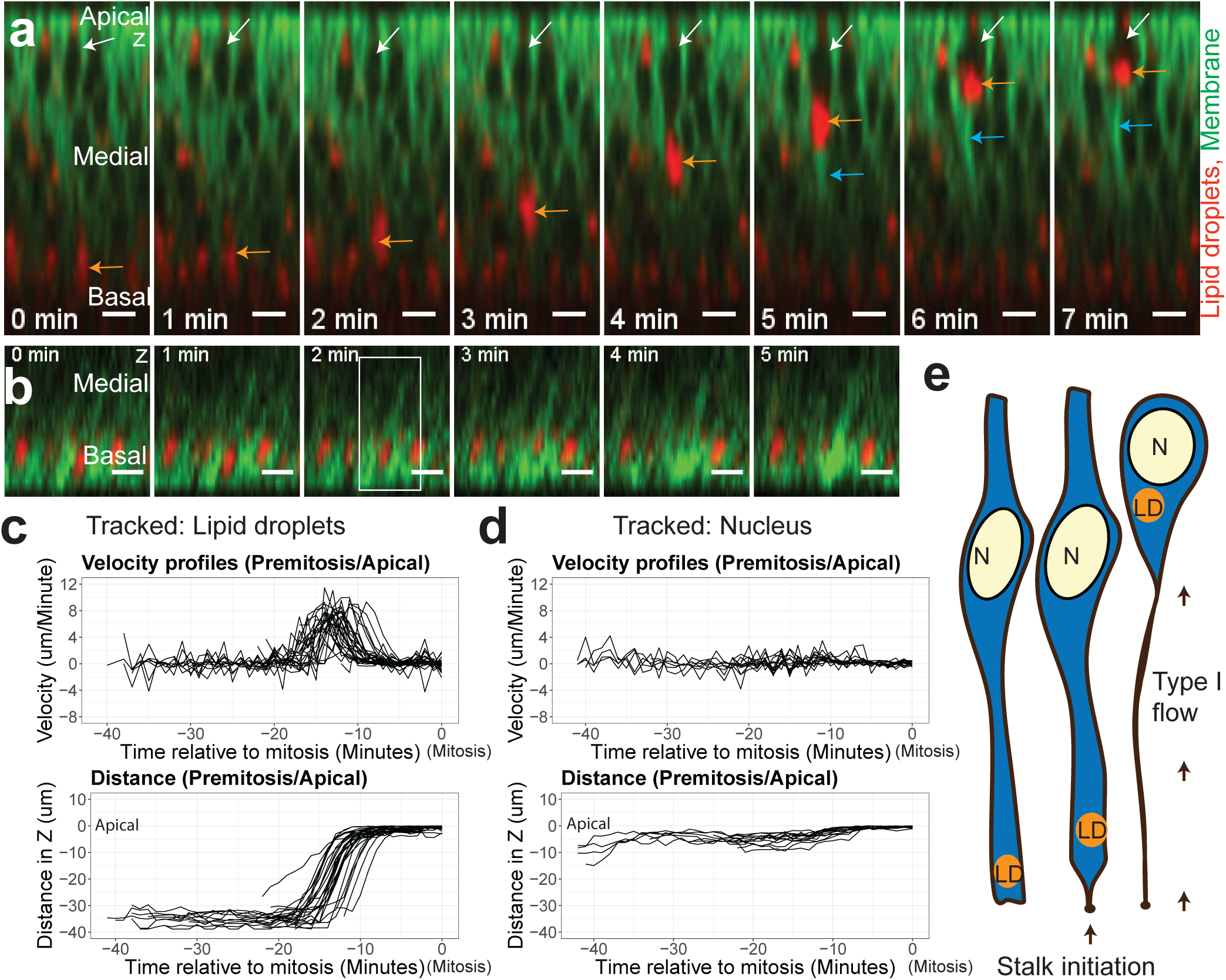
A basal-apical (type I) flow displaces the lipid droplets and nucleus to apical surface with a stalk initiated at basal surface. (a, b) *c765-GAL4>UAS-PLCdelta-PH-GFP* with 1:2000 LipidTOX Deep Red. (a) Representative images of apical-directed type I flow preceding mitosis of lipid droplet clusters (LDC, orange arrow), nucleus (white arrow), and stalk (blue arrow). (b) z slice of time series of the stalk formation at the basal surface. Arrows depicts process formation and square represents the first time point where the stalk was first visible. (c) Velocity and distance profiles tracked LDC in the apical-basal axis (N = 29 LDC). Distance values are normalized to mitosis time point (x axis = 0) and apical surface (y axis =0). (d) Velocity and distance profiles tracked LDC in the apical-basal axis (N= 12 nuclei). Distance values are normalized to mitosis time point (x axis = 0) and apical surface (y axis =0). (e) Model representing the type I flow. Scale bars: 5 µm.

LDC trajectories associated with this type I flow initially showed large periods of little movement at the basal surface (resting motion). This was followed by an interval of apical-directed motion, defined as time window where the LDC attains a minimum velocity of 1 µm/minute, maintains a positive velocity for at least 5 minutes and ends if the velocity drops below 0.1 µm/minute). This was followed by a short time period with little movement until the end of mitosis (when cytokinesis cleaves the mother cell; Supplementary Fig. S2a). The Interval of apical-directed motion lasted on average (9.65 ± 2.52 minutes, mean ± SD, N = 29) and LDC associated with it had an average velocity of (3.41 ± 2.88 µm/minute, mean ± SD, N = 29). Mitosis occurred on average (7.27 ± 1.84 minutes, mean ± SD, N=29) from the end of the Interval of apical-directed motion. It should be noted that the nucleus in these mitotic cells was already close to the apical surface as previously observed (Meyer et al., 2011) and nuclear displacement in type I flows was less extensive than the movement of lipid droplets (Fig. 2c-d, Supplementary Fig. S2c). The observed flow was associated with a bright basal process connecting to the basal surface that was previously shown to be highly enriched in actin (Meyer et al., 2011). We refer to this basal process as the basal stalk. Consistent with the finding that high levels of PIP_2_ are often associated with actin polymerization (Logan and Mandato, 2006), PLCδ PH-GFP was found to be highly enriched on this basal stalk (Fig. 2a-2b). The basal stalk formed within a time period of maximally one minute (N=6) and displaces the LDC apically (Fig. 2b). Type I flows were also observed during the live imaging of a mitochondrial marker (Supplementary Fig. S2d).

### Type II apical-directed flow moves the nucleus to apical-medial position

We found evidence for a second type of apical-directed flow for LDC in cells where the nucleus was located closer to basal surface (Fig. 3a-c, Supplementary Mov S3). We define this flow as type II flow. While type I flows transported the LDC to the mitotic zone, type II flow transported the nucleus to a more apical-medial position (Fig. 3c). The motion of the LDC and the nucleus was correlated for periods of apical-directed motion (Pearson coefficient = 0.83 ± 0.14, mean ± SD, N=10 LDC and nucleus pairs). The dynamics of type II flow (Fig. 3b) showed slower velocities and longer time scales than type I flow (Fig. 2c). If only apical-directed movements (positive velocities) were considered, the average velocity of LDC was (0.78 ± 0.84 µm/minute, mean ± SD, N = 10). We did not observe mitosis during the imaged time periods. In one cell, the LDC were observed to be moving towards basal surface after the nucleus reached its apical position.

**Fig 3:**
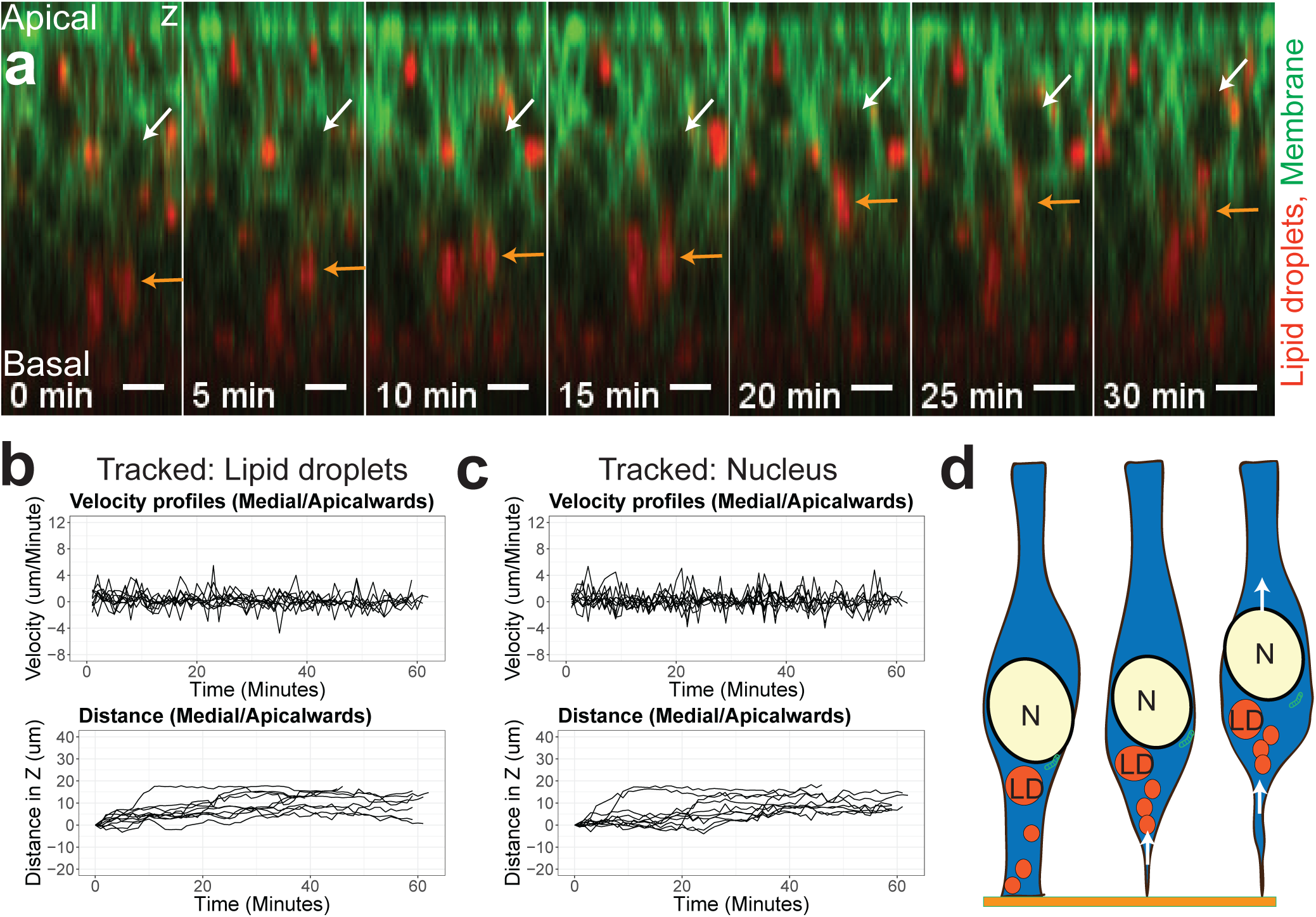
A basal-apical flow (type II) flow displaces the lipid droplets and nucleus from a basal-medial to an apical-medial position. (a) *c765-GAL4>UAS-PLCdelta-PH-GFP* with 1:2000 LipidTOX Deep Red. (a) Representative images for Type II flow of LDC from basal surface that leads to an apical-directed motion of the nucleus. Lipid droplet clusters (LDC, orange arrow), nucleus (white arrow). (b) Quantification of velocity and distance profiles of tracked LDC in the apical-basal axis (N = 10 LDC). (c) Quantification of velocity and distance profiles of tracked nuclei in the apical-basal axis (n=10 nuclei). (d) Model of Type II flow. Scale bars: 5 µm.

### Basal directed flow and ground state of LDC motion

Post mitosis, we noticed a basal-directed flow of LDC accompanying the nucleus. (Supplementary Fig. S3a-a’’, Supplementary Movie. S3). For measuring the ground state of LDC motion, we imaged the basal surface of regions that did not undergo mitosis (Supplementary Fig. S3b-b’, Supplementary Movie. S3).

### Actin perturbation with Latrunculin B treatment

Since the nuclear motion was the result of the actin-myosin contractility (Meyer et al., 2011; Norden et al., 2009), disruption of actin polymers should affect the cytoplasmic flow. We employed Latrunculin B (LatB) to inhibit the polymerization of actin (Wakatsuki et al., 2001). We noticed that the mitotic events were blocked, no directed motion of LDC across the cell was observed indicating both type I and type II LDC flows ceased (Fig. 4a-a’, Fig. 4b-b’, Supplementary Movie. S4).

**Fig 4:**
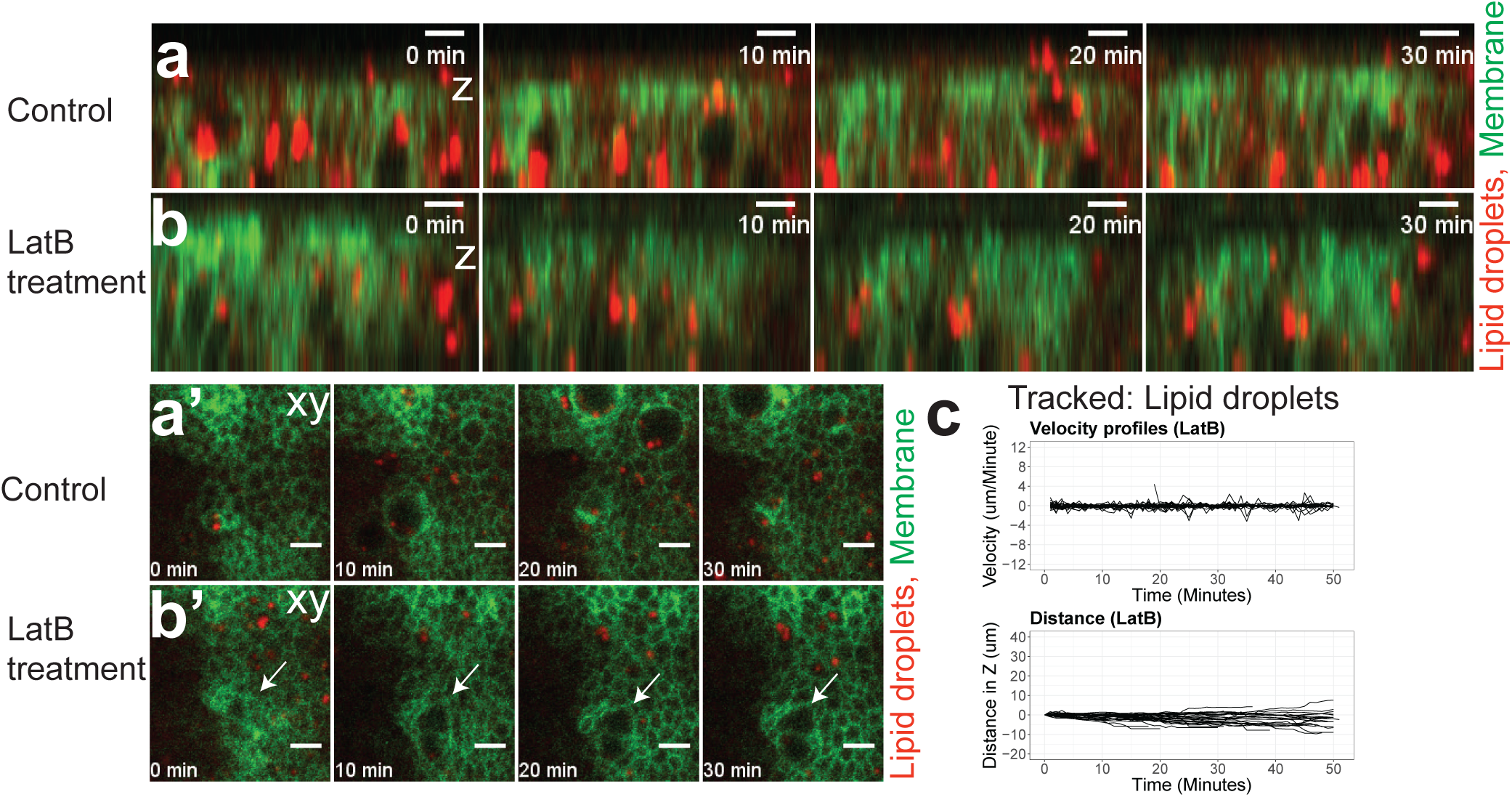
Targeting actin with Latrunculin B (LatB) treatment switches off the flows. (a, a’, b, b’) *c765-GAL4>UAS-PLCdelta-PH-GFP* with 1:2000 LipidTOX Deep Red. (a, b) z slice. (a’, b’) xy slice. (a, a’) Time series Control. (b, b’) Treatment with 50 µM LatB. (b’) Cell blocked in late telophase (white arrow). (c) Quantification of velocity and distance profiles of tracked LDC in treated discs (n=35 LDC). Scale bars: 5 µm.

### Mean square displacement and periodogram analysis

Mean square displacement (MSD) analysis was commonly used to describe whether the mode of displacement particles is stochastic/similar to free diffusion or advection/directed transport. It had been previously used to study the stochastic and directed properties of nuclei during IKNM (Norden et al., 2009). MSD curve of type I LDC flow is advection (Supplementary Fig. S4a). MSD curves of type I nucleus flow, post mitotic basal-directed flow and the type II flows (Supplementary Fig. S4b) have characteristics of advection. MSD curves of ground state of LDC (Supplementary Fig. S4c) and LatB treated LDC showed that the motion is stochastic.

A periodogram calculates the dominant frequencies of a time series to identify periodic events. Average periodogram of type I LDC flow contained predominately low frequency components while the other flows were mixed with some high frequency components (Supplementary Fig. S4e-g). Average periodogram of LatB treated LDC revealed a reduction of all frequencies compared to ground state of LDCs (Supplementary Fig. S4g-h).

### Viscosity of the cytoplasm

The diffusion coefficient of this ground state of LDC was calculated as (D = 0.0027 ± 0.0024 µm^2^/sec, mean ± SD, N = 27) by a linear fit of the MSD curves (Tarantino et al., 2014). Using the Stokes–Einstein equation (Supplementary Eq. 1) with *D* = 0.0027 µm^2^/sec, T = *298* K, mean radius of lipid droplets r = *0.64* µm, the dynamic viscosity of the cytoplasm was 0.126 Pa*sec or about 126 times more viscous than water (0.001 Pa*sec).

### Aspect ratio of packing of LDC

A LDC contains multiple lipid droplets that were often closely packed that they could not be distinguished individually by light microscopy. Aspect ratio of the LDC was calculated by fitting an ellipse to the segmented LDC. We noticed that LDC exhibited different aspect ratios between the type I (Fig 5a-a’) and type II (Fig 5b-b’) flows implying different packing configurations of the lipid droplets. We noticed that type I LDC was tightly packed with low aspect ratios (circle) in the apical-basal axis while Type II LDC signal was more loosely packed during some time periods (ellipse) and tightly packed (closer to circle) during other time points. The aspect ratio of type II LDC (Fig. 5a-a’) (5.21 ± 1.88, mean ± SD, N=59) is larger compared to the aspect ratio of type I LDC (Fig. 5b-b’) (2.03 ± 0.9, mean ± SD, N=23) indicating a tighter packing of lipid droplets during Type I flow (Fig5C). The average area of Type I lipid droplets (31.85 ± 6.11 µm^2^, mean ± SD, N=23) is much larger compared to average area of Type II lipid droplets (11.46 ± 3.66 µm^2^, mean ± SD, N=59).

**Fig 5:**
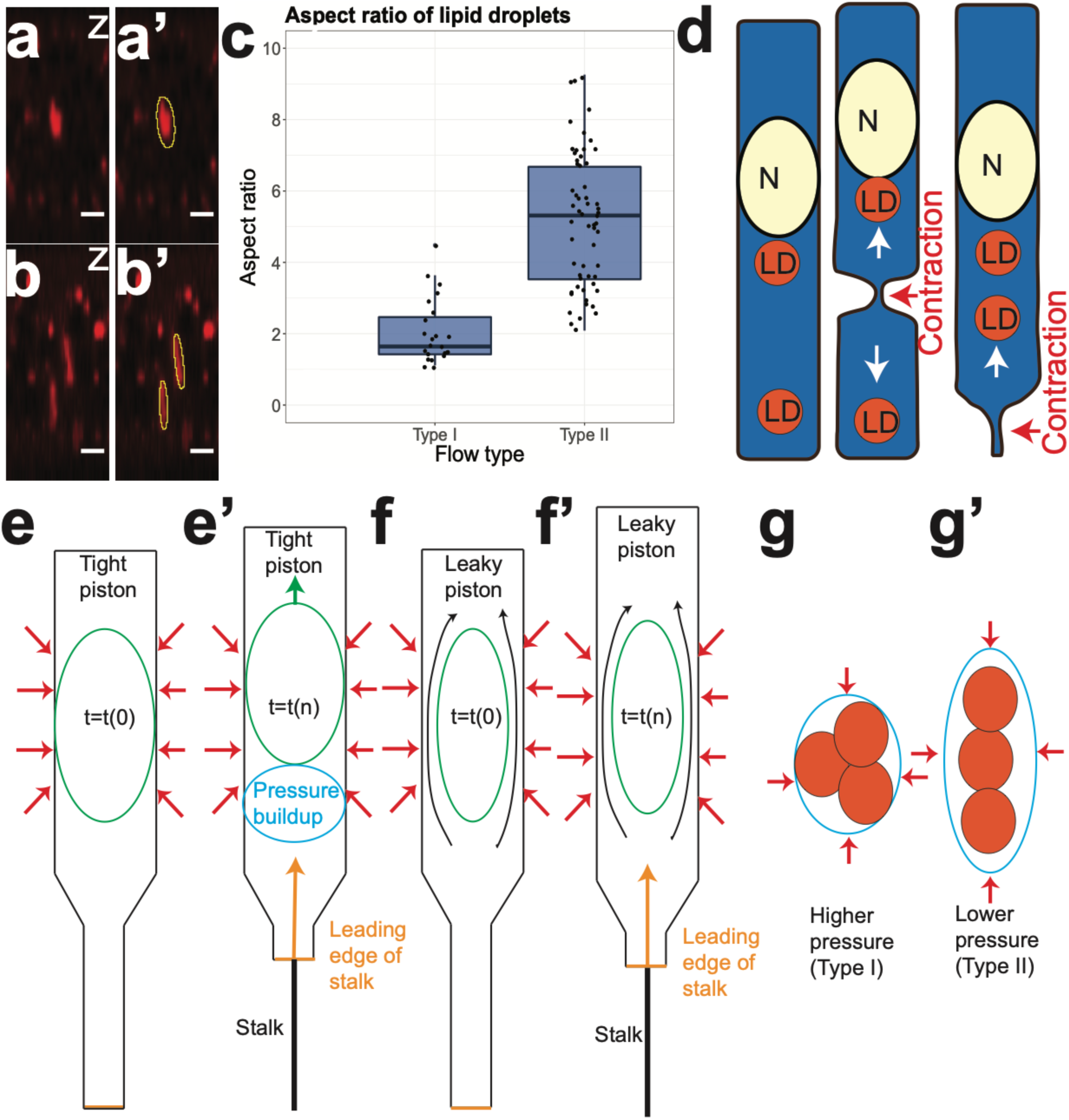
Aspect ratio of LDC and nuclear piston. (a-a’) Representative images of LDC in Type I flow and ellipse fitting. (b-b’) Representative images of LDC in Type II flow and ellipse fitting. (c) Quantification of aspect ratio (major/minor axis lengths of ellipse that was fitted to the LDC). (d) Model representing the pinching/squeezing force and generation of flows of LDC. In model (e-e’) the nucleus acts as a tight seal prevent leaking of cytoplasm. The pressure built up under the nucleus will propel the nucleus up along with the basal process. In model (f-f’) the nucleus acts as a leaky seal where cytoplasm can flow around the nucleus. Red arrows represent the compression through force from neighboring cells. (g) Model representing relationship between aspect ratio and pressure in type I flow. (g’) Model representing relationship between aspect ratio and pressure in type II flow. Scale bars: 5 µm.

### Low Reynolds number and domination of viscous forces

Reynolds number is a dimensional value that is used to measure the ratio of inertial forces to viscous forces. For very small Reynolds numbers at the cellular scales, viscous forces were known to dominate (Purcell, 1977). The Reynolds number for a flow (Supplementary Eq. 4) in a tube was calculated as 1.421*10^−6^ for type I and 3.25*10^−7^ for type II flow. ρ = 1050 kg/m^3^ ((Handel, 2017)), *ϑ* = 3.41 µm/minute as mean velocity for type I Interval of apical-directed motion and *ϑ* = 0.78 µm/minute as mean velocity for positive type II flow, D= 50 µm and μ = 0.126 Pa*s.

### Inverse relationship between particle size and velocity

The velocity differences of the type I and type II flows may be accounted by the differences in size of the transported cargo. An inverse relationship between velocity and particle size could be derived for 1D motion from the Stokes–Einstein equation and the MSD equation for mobility analysis (Supplementary Eq. 3). Nuclei and lipid droplets were 3D segmented to calculate volumes. The type I flow was based on the motion of lipid droplets while the type II flow was based on the motion of both nucleus and lipid droplets. Since size of the nucleus (Volume = 56.49 ± 12.05 µm^3^, mean ± SD, N= 112, mean ± SD, N=112) was much larger than a lipid droplet (Volume = 3.02 ± 2.77 µm^3^, mean ± SD, N=2573), we bundled the nucleus and LDC together as one particle with the combined radius for the type II flow for this analysis. The mean radius *r*_*N*_ of the nucleus was 2.16 µm and the mean radius *r*_*N*_ of lipid droplets was 0.64 µm. Applying these values to the ratio of radii

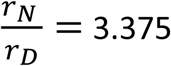

The mean velocity of the Interval of apical-directed motion for type I LDC *v*_*D*_ was 3.41 µm/minute. The mean velocity of positive type II flow, *v*_*N*_ of the nucleus was 0.78 µm/minute. Applying these values to the velocity ratio,

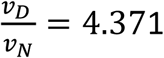

The similar values of these inverse relationship between radius and velocity suggest that the size of the particles may be one factor for the velocity differences between type I and type II flows.

### Volume of mitotic cells

Since most of the cytoplasm was at the apical surface, it was possible to estimate the volume of cells by 3D segmentation in different phases of mitosis (Supplementary Fig. S6b). Although it was difficult to optically segment the stalk, we calculated its approximate volume to be (19.77 ± 13.21 µm^3^, mean ± SD, N=210) by interpolating the last segmented area of the stalk. Finally, we examined sibling daughter cells in telophase and noticed that their volumes and cytoplasmic components were not always equally distributed (Supplementary Fig. S6a’-a’’).While the volume difference was minimal (7.5 ± 2.3 % of combined volume, mean ± SD), the lipid droplet difference seemed to be more substantial (larger cells (9.11 ± 3.55 µm3, mean ± SD), smaller cells (4.66 ± 2.44 µm3, mean ± SD).

## Discussion

Previous studies on IKNM have focused on the migration of the nucleus from the medial zone into the mitotic zone prior to migration and, following mitosis, a slower migration of nucleus back into the medial zone driven by acto-myosin contractile ring (Meyer et al., 2011; Norden et al., 2009; Sauer, 1935). However an acto-myosin ring undergoing contraction would displace the highly viscous cytoplasm (126 times more viscous than water) generating a flow that would displace cytoplasmic components such as lipid droplets and mitochondria. To demonstrate the cytoplasmic flows, we tracked the lipid droplets and identified two types of the apical-directed IKNM and one type of basal-directed IKNM.

### Properties of cytoplasmic flows

Prior to mitosis, the apical-directed type I flow pushes cytoplasm containing lipid droplets into the mitotic zone (Fig. 2e). Following mitosis, cytoplasm with lipid droplets was transported towards the basal surface by a basal-directed flow (Supplementary Fig. S3a’’). at a much slower rate, this flow has the similar characteristics of the basal-directed nucleus migration following mitosis (Sauer, 1935). The apical-directed type II flow (Fig. 3d) pushes basal-medial nuclei to a more apical-medial plane. Though we have not observed subsequent cell divisions associated with the type II flow, we cannot exclude the possibility that these occur at time scales that exceed our imaging periods. If the order of the flows needs to be considered in the context of traditional IKNM system where more basal nuclei are further in cell cycle (Sauer, 1935), then type II flow would precede the type I flow by displacing the nucleus to a much more apical plane. Type I flows were associated with larger and tightly packed LDC compared to type II flows. Type I flows were faster in velocity, spread over shorter time scales and directed from basal to apical surface. Type II flows were slower in velocity spread over longer time scales and were mixed with time periods of stochastic behavior. Type I flow were associated with the pronounced basal stalks which can provide both directionality and a tight seal by preventing flow in the opposite direction. Neither the slower basal-directed flow nor the slow apical-directed type II flow was accompanied with an apical or basal stalk enriched with actin and PIP_2_, respectively. All the cytoplasmic flows ceased when cells were treated with LatB, an actin inhibitor. This showed that these flows were driven by a contractile acto-myosin ring similar to what has been previously proposed with nuclear motion (Mayer et al., 2010; Meyer et al., 2011).

### Shape of the nucleus as indicator of forces and direction of motion

Already 80 years ago (Sauer, 1935), the pointed droplet-like morphology of the nucleus during IKNM was described. Recent studies have proposed that nuclear shape changes might serve as an indicator for the direction of motion and forces acting on the nucleus (Lee and Norden, 2013; Zhao et al., 2012). In the TEM data, we observed droplet shape of nuclei with a close proximity to clusters of lipid droplets and mitochondria showed evidence of the apical directed type I (Supplementary Fig. S5d), type II flows (Supplementary Fig. S5b-c) and basal directed flow (Supplementary Fig. S5f).

### Nuclear piston and increased hydrostatic pressure generates the lifting force

It had been demonstrated that nucleus can act as a piston to compartmentalize the pressure where an increased hydrostatic pressure between the nucleus and leading edge of the cell drives cell migration (Petrie et al., 2014). Models, in which the nucleus acts as a piston in a hydraulic cylinder (cell), could be applied in IKNM to explain the mechanism behind nuclear migration as an effect of the flows. When the contractile machinery (Fig. 5d) generates a force displacing the cytoplasm towards the nucleus, the nucleus could either function as a tight (Fig. 5e-e’) or as a leaky piston (Fig. 5f-f’). In the tight piston model (Fig. 5e), it is assumed that the cell and its nucleus are under compression from neighboring cells forming a tight seal and preventing cytoplasm from passing around it. The leading edge of the stalk pushes the cytoplasm towards the nucleus, this directly increases the hydrostatic pressure under the nucleus until at some point the force suffices to push the nucleus up (Fig. 5e’). In the leaky piston model (Fig. 5f), the nucleus does not form a tight seal with the surrounding cell membrane. So when the leading edge of the stalk or contractile ring pushes cytoplasm towards the nucleus, some cytoplasm bypasses the nucleus and does not allow for accumulation of hydrostatic pressure under the nucleus. The nucleus only moves when it is in direct contact or in close proximity to the leading edge of the stalk or contractile ring and thus exposed to a direct force from it (Fig. 5f’). Evidence of the effect of the hydrostatic pressure could be seen in three types of data. First, in TEM data for cells being in the type I and type II flow (showing droplet like morphology of the nucleus and close proximity of nucleus to LDC) and where lipid droplets were almost touching the nucleus, the nuclear envelope showed a slight deformation (Supplementary Fig. S5b, d). Second, In TEM data for nuclei which fit the criteria of being in a flow, we did not observe any lipid droplets in the gap between the nucleus and the cell membrane. It could be possible that lipid droplets are too large to fit into this gap since the average diameter of lipid droplets was (1.82 ± 0.42 µm, mean ± SD, N=167) and the gap between nuclei and cell membranes was (0.122 ± 0.04 µm, mean ± SD, N=356). We confirmed that lipid droplets do not move around the nucleus during Type I or Type II flow from the live imaging data. Third, we noticed from live imaging of type I flows that the smaller mitochondria, embedded in the cytoplasm, tend to move with the same motion and don’t move around the nucleus (Supplementary Fig. S2d, Supplementary Movie. S3). From these observations it is likely that the nucleus forms a tight seal that prevents leakage due to hydrostatic pressure on the nucleus.

### Post mitotic distribution of cytoplasm and lipid droplets

Post mitosis, the cytoplasmic volume of the mother cell was not equally distributed to the two daughter cells, further the larger daughter received more lipid droplets. This may result in a potential growth advantage over the smaller sibling, as lipid droplets are known to serve as energy storage (Beller et al., 2010; Grönke et al., 2003). It had been observed previously that cells in the wing disc do not pass through the cell cycle in clusters and cell division rate is not clonal (Milán et al., 1996). Volume and resource differences may have an influence in desynchronizing cell cycle progression of sibling daughter cells, and thus on the stochastic nature of the cell division pattern in the wing imaginal disc.

In conclusion, by tracking and measuring the dynamics of lipid droplets during IKNM, we demonstrated that cytoplasmic flows were responsible for nuclear migration by increasing the hydrostatic pressure under the nucleus which functions like a piston. We have identified the existence of a type I flow preceding mitosis, a type II flow for medial nuclei and a basal flow in post mitotic cells. Treatment of the wing disc with Latrunculin B ceased the flows, showing that they are driven by acto-myosin contractile forces. Future research will establish whether the phenomenon and the function of cytoplasmic flows during IKNM are conserved across species.

## Methods

### Fly stocks and genetics

The following stocks were used: *y w ubi-GFP[nls] hsp-flp* FRT19, *P{w[+mC]=*UAS-*mito-HA-GFP.AP}3*, e[1] (Bloomington 8443), w[*]; *P{w[+mC]=4C.*UAS*-PLCdelta-PH-mCherry}CU166* (Bloomington 51658), *y[1] w[*]; L[1]/CyO; P{w[+mC]=*UAS-*PLCdelta-PH-EGFP}3/TM6B, Tb* (Bloomington 39693), *ap-*GAL4 tub-GAL80^ts^*/ CyO; Mkrs/ TM6B, Sp/ CyO;c-765-*GAL4*/ TM6B.* For UAS-GAL4 tub-GAL80^ts^ experiments, larvae were kept at 29c for 48 hours

### Sample preparation

For living imaging, wing discs were mounted in an imaging chamber with a filter on top as described in (Zartman et al., 2013). For fixed imaging, samples were dissected in Ringers medium, were kept at 4% PFA overnight at 4°C before Immunostaining. Wing discs were mounted in VECTASHIELD (Vector Labs) inside a small cavity formed with reinforcement rings attached to the slide and avoiding compression from the cover slip.

### Immunofluorescence

1:2000 LipidTOX Red Neutral Lipid Stain (Thermofisher). 1:2000 HCS LipidTOX Deep Red neutral lipid stain (Thermofisher). 1:400 of 1mg/ml PH3 (Millipore) antibody.

### Drug treatments

20 mM stock solution of Latrunculin B in DMSO was diluted 1:400 in Schneider’s medium to a working concentration of 50 µM.

### Ultrastructure analysis of wing discs

Chemical fixation: Wing discs were dissected from L3 larvae in Schneider’s medium and transferred onto 6 mm sapphire discs. Wing discs were fixed in 2.5 % glutaraldehyde for 6 hours at room temperature, rinsed with 0.1 M cacodylate buffer, incubated in 1 % osmium tetroxide in 0.1 M cacodylate buffer at room temperature followed by rinsing with pure water. Subsequently, samples were kept at 4°C for 1.5 hours in 1 % uranyl-acetate in water, followed by a water rinsing step and dehydration with a graded alcohol series (70 % for 20 minutes, 80 % for 20 minutes, 96 % for 45 minutes, 100 % for 15 minutes). Samples were then transferred to 100% propylene oxide for 30 minutes, incubated in 50% Epon/Araldite in propylene oxide at 4°C for two hours, and in 100% Epon/Araldite in a flat silicon embedding mold for one hour before polymerization at 60°C for 24 hours.

Ultra-thin sections were contrasted with Reynolds lead citrate and imaged in a CM 100 (at 80 kV) or Tecnai Spirit G2 (at 120 kV) transmission electron microscope (Thermo Fisher Scientific, Eindhoven, The Netherlands) using a side-mounted Gatan Orius 1000 digital camera and the Digital Micrograph software package (Gatan, Munich, Germany).

### TEM alignment

TrakEM2 (Cardona et al., 2012) was used to align the TEM multiple tiles with least squares montage mode followed by elastic montage mode.

### Confocal Imaging

Confocal images were acquired on a Zeiss LSM710. For fixed samples, confocal stacks were acquired with 63x 1.4 NA oil immersion objective with settings for deconvolution. For larger overviews, 40x 1.3 NA (oil), or 25x (oil) objectives were used. For live samples, confocal stacks were acquired with 40x 1.2 NA water immersion objective at 1 minute intervals. Three wing discs were mounted apical side closer to the objective, imaged sequentially at 1 minute intervals. For imaging basal side for visualization of stalk formation, discs were mounted basal side closer to the objective.

### Image analysis

Average periodograms were calculated using the Genecycle (Ahdesmaki et al., 2012) package in R (R Core Team, 2016). Correlation coefficients were calculated with “*cor”* function from stats package in R. Cross correlations were calculated with “*ccf”* function from stats package in R. MSD analysis was performed with class MSDanalyzer (Tarantino et al., 2014) in MATLAB R2016B (MathWorks). Images were deconvolved with Huygens Professional version 17.04 (Scientific Volume Imaging, The Netherlands,<u>__</u>http://svi.nl__).

### Tracking LDC and nuclei

Areas around mitotic cells were taken as regions of interest and sectioned in z. In the z sections, LDC and nuclei were detected with LOG detector and tracked with trackmate (Tinevez et al., 2017) plugin in Fiji (Schindelin et al., 2012). Manual segmentation was made for missing particles using other slices as reference and filters (x, y, quality) were applied to restrict detection to the region of interests around the nuclei. LDC were detected with 3 - 5 micron blob diameter, 0 - 40 threshold and median filter settings. The low intensity regions in the z sections of cell membrane marker corresponded to nuclei as seen in the example with lamin-GFP nuclear marker (Supplementary Fig. S2b-b’). For nuclei detection, we tracked inverted signal of the cell membrane marker. Nuclei were detected with 7 - 10 micron blob diameter, 0 - 40 threshold and median filter settings. The y coordinates and time of the tracks were exported to csv files. The plotting and analysis was continued in R.

Mean, median, variance and SD were calculated separately for each of the tracks. The global means and variances were then computed from those means and SD were computed using “*combinevar*” function from *fishmethods* package (Nelson, 2017) in R (R Core Team, 2016). It should be noted that we used the complete type I LDC trajectory for the MSD, frequency calculations without exclusions for consistency, unless indicated specifically by the term interval of apical-directed motion.

### Cell volume analysis

Cells were identified in different stages of mitosis (prophase, metaphase, anaphase, telophase, and cytokinesis) from the morphology of nucleus in DAPI staining in the mitotic zone with a terminology similar to (Meyer et al., 2011). Cell shape in 3D was segmented using ilastik (Sommer et al., 2011) based on PH-EGFP. BoneJ plugin (Doube et al., 2010) in Fiji (Schindelin et al., 2012) (Java 6 life line version) was used to calculate volumes from the segmented masks. In Fiji, masks (3x) from ilastik were diluted and then applied on the images to reduce the amount of background.

### Lipid droplet, nuclear volume analysis

Lipid droplet masks were calculated by median filter of 0.5, Otsu (OTSU) threshold in Fiji. *Lamin*-GFP was used to segment nuclei separated by with nuclear distance to restrict merging of meshes in ilastik (Sommer et al., 2011). Volumes are calculated by BoneJ plugin (Doube et al., 2010) in Fiji (Java 6 life line version). A size limit was imposed on nuclear sizes with an upper limit of 80 um^3^, masks above this limit were observed as segmentation artifacts and excluded from the analysis. Radii of lipid droplet (0.84 ± 0.2 µm, mean ± SD, N=2573) and nucleus (2.36 ± 0.17 µm, mean ± SD, N=112) were calculated assuming the particles were spheres. The upper limit of axial resolution that can be resolved from confocal microscopes is in the order of 100 nm (Schrader et al., 1996). Due to the high levels of photon noise associated with fluorescence images (Huygens de-convolution FAQ), we used a conservative estimate of 200 nm as the axial resolution. After subtracting this axial resolution, the radii of lipid droplets (0.64 ± 0.2 µm, mean ± SD, N=2573) and nucleus (2.16 ± 0.17 µm, mean ± SD, N=112).

### Aspect ratio/ellipse fitting of LDC

A median filter of 0.5 was applied on the LDCs in the lateral sections, thresholded according to Otsu and ellipse fitting was made with the *“Analyze Particles”* plugin in Fiji on the ROIs.

### Lipid droplet counting in cells

Cell shape in the wing pouch was segmented with find maxima plugin in Fiji. These segmented cell shapes were overlaid on the channel with the lipid droplets, detection of number of lipid droplets was made with the *“Find Maxima”* plugin in Fiji. Average of lipid droplets per cell was the number of segmented cells divided by the number of droplets detected in those segmented cells. We divided the overall analysis into two types. Fine analysis (Supplementary Fig. S1c) was focused in the mitotic zone with the cells classified into dividing and non-dividing cells according to mitotic rounding. Coarser analysis (Supplementary Fig. S1c’) was made from apical to basal surface. For finer analysis, the lipid droplets detected were (23 ± 14, mean ± SD) and cortices detected were (1827 ± 277, mean ± SD).For coarser analysis, the lipid droplets detected were (461 ± 300, mean ± SD) and cortices were (2278 ± 178, mean ± SD).

### Volume of the stalk

The volume of the stalk was calculated by multiplying the segmented area of the thinnest regions of the stalk (just before it was no longer possible to differentiate it from the surrounding cells) and multiplying it with the height (the end of mitotic zone to the basal surface).

## Author contributions

AB conceived the project, performed the experiments and data analysis and wrote the manuscript. AK performed the ultrastructure preparation for TEM imaging.

## Acknowledgments

I would like to thank Konrad Basler for the position in his laboratory and support as my PhD advisor. I would like to thank Simon Tanaka, Robert Witte and Dominik Eder for their valuable inputs, feedback and discussion during the writing of this manuscript. I would like to thank Bloomington Drosophila Stock Center for their fly stocks. I would like to thank Carmen Kaiser for TEM preparation. I would like to thank Center for Microscopy and Image Analysis, University of Zurich for their TEM equipment. This work was supported by the University of Zurich.

