## Supplementary Data for "Cytoplasmic flows caused by actomyosin contraction drive interkinetic nuclear migration"

**Mobility analysis**

Stokes–Einstein equation for diffusion of spherical particles through a liquid$D = \frac{k_B T}{6\pi\eta r}$ (1a)

$D$ is diffusion coefficient, $k_B$ is Boltzmann's constant, $T$ is absolute temperature, $\eta$ is dynamic viscosity, $r$ is radius of the spherical particle.

$$D \propto 1/r \tag{1b}$$

For 1D motion of a spherical particle, the relationship between mean-squared displacement and diffusion coefficient is

$$(\Delta x)^2 = Dt \tag{2a}$$

$(\Delta x)^2$ is mean-square displacement, $D$ is diffusion coefficient and $t$ is time.

$$v \propto D \tag{2b}$$

$v$ is velocity and $D$ is diffusion coefficient

Combining (2A and 2B) and assuming lipid droplets and nucleus are spherical particles. radius $r_N$ of the nucleus, radius $r_D$ of the lipid droplets, average velocity of the type I LDC flow, $v_D$ and the average velocity $v_N$ of type II nucleus.

$$\frac{r_D}{r_N} = \frac{v_N}{v_D} \tag{3}$$

**Reynolds number for a flow in a tube**

$$R_e = \frac{\rho \cdot v \cdot D}{\mu} \tag{4}$$

$R_e$ is Reynolds number (dimensionless), ρ is Density of the fluid (kg/m$^3$), $v$ mean velocity of the fluid (m/s), D is tube diameter (m) and μ is dynamic viscosity of the fluid (Pa*s).

### Supplementary Figures

Fig S1

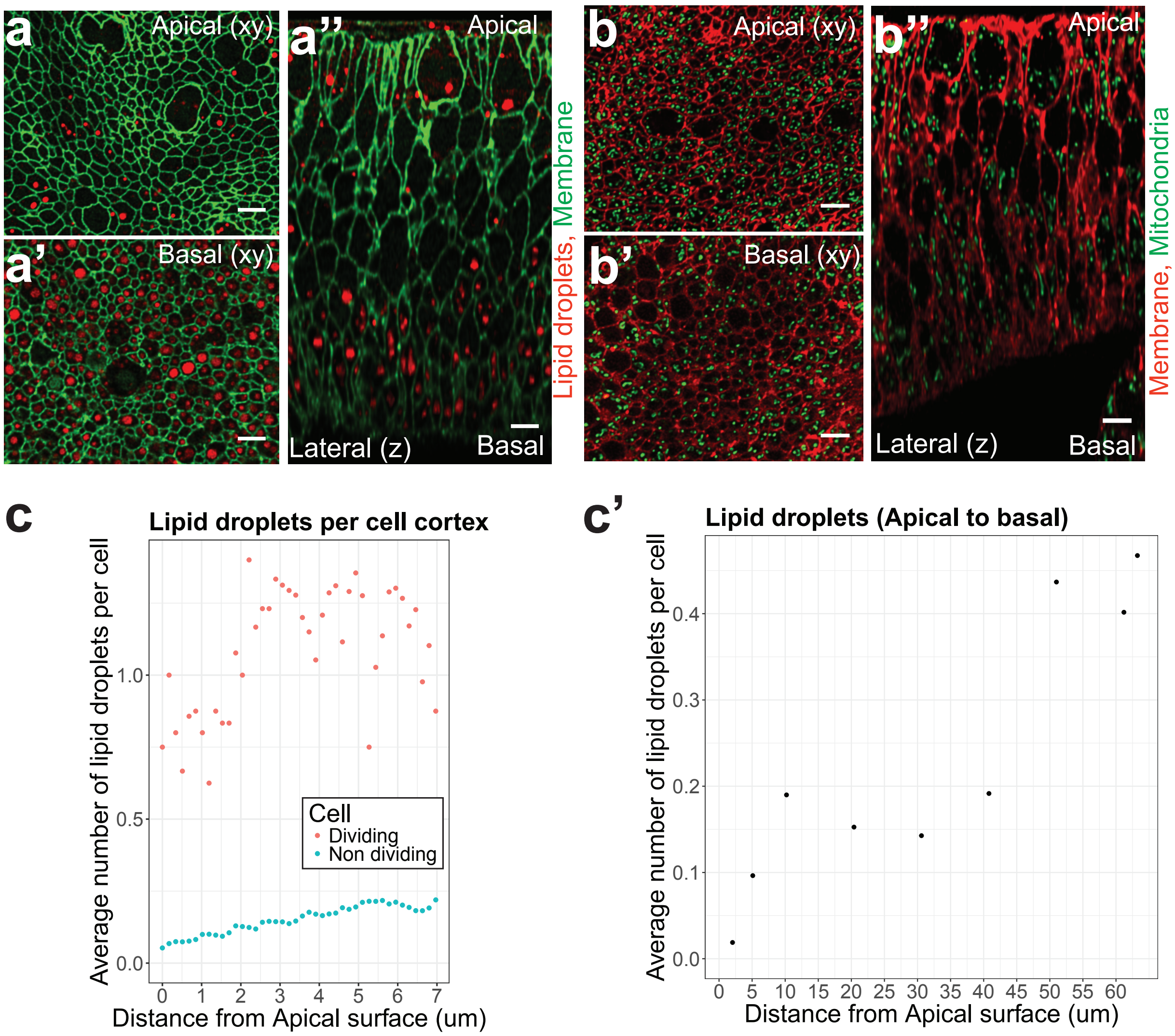


Fig S1: Spatial distribution of lipid droplets and mitochondria.

(a-a'') *c765*-GAL4>UAS-*PLCdelta-PH-GFP* with 1:2000 LipidTOX Deep Red. Representative images of lipid droplets distribution in apical, basal and lateral sections. (b-b'') *ap*-Gal4 tub-GAL80$^{ts}$; UAS-*mito-HA-GFP*>UAS-*PLCdelta-PH-*

*mcherry.* Representative images of mitochondria distribution in apical, basal and lateral sections. (c) Quantification of average LDs per cell in the mitotic zone. Images were quantified with z spacing of 0.17 µm. (c') Quantification of average LDs per cell for all the pouch cells from apical to basal surface with a z spacing of 5 µm. Scale bars: 5 µm.

Fig S2

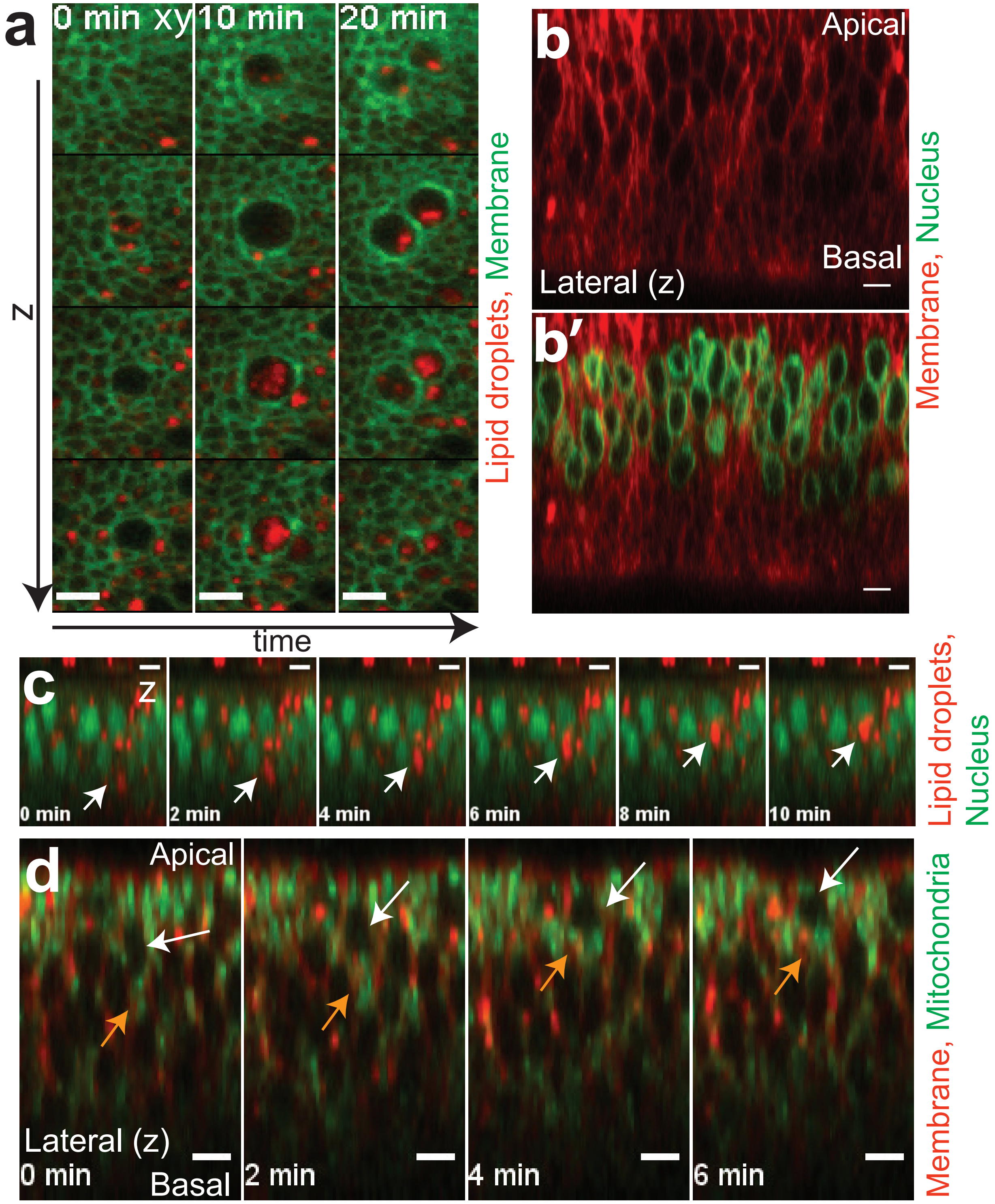
a
0 min xy
10 min
20 min
z
time
Lipid droplets, Membrane
b
Apical
Basal
Lateral (z)
b′
Membrane, Nucleus
c
z
0 min
2 min
4 min
6 min
8 min
10 min
Lipid droplets, Nucleus
d
Apical
Lateral (z)
0 min
Basal
2 min
4 min
6 min
Membrane, Mitochondria

Fig S2: Type I flow in xy/z slices and mitosis, distribution of nuclei in lateral view, type I flow with nuclear marker, and type I flow with mitochondria. (a) *c765-*GAL4>UAS-*PLCdelta-PH-GFP* with 1:2000 LipidTOX Deep Red. Time-lapse confocal microscopy imaging of multiple Z sections at apical surface spaced at 2.5 µm to illustrate the appearance of lipid droplets in the time preceding mitosis. (b-b') *ap-*Gal4 tub-GAL80$^{ts}$; UAS-*lam -GFP*>UAS-*PLCdelta-PH-mcherry.* (b) Low intensity regions in the lateral section of the tissue represents nucleus as observed in the lamin nuclear marker in (b') . (c) ubi-nlsGFP with 1:2000 LipidTOX Deep Red. Time series of lateral view representing the nuclear migration during type I flow. LDC is represented by white arrow (d-d') *ap*-Gal4 tub-GAL80$^{ts}$; UAS-*mito-HA-GFP*>UAS-*PLCdelta-PH-mcherry.* Time series of lateral view cytoplasmic motion with mitochondrial (orange arrow) marker during type I flow and the nucleus (white arrow). Scale bar: 5 µm.

Fig S3

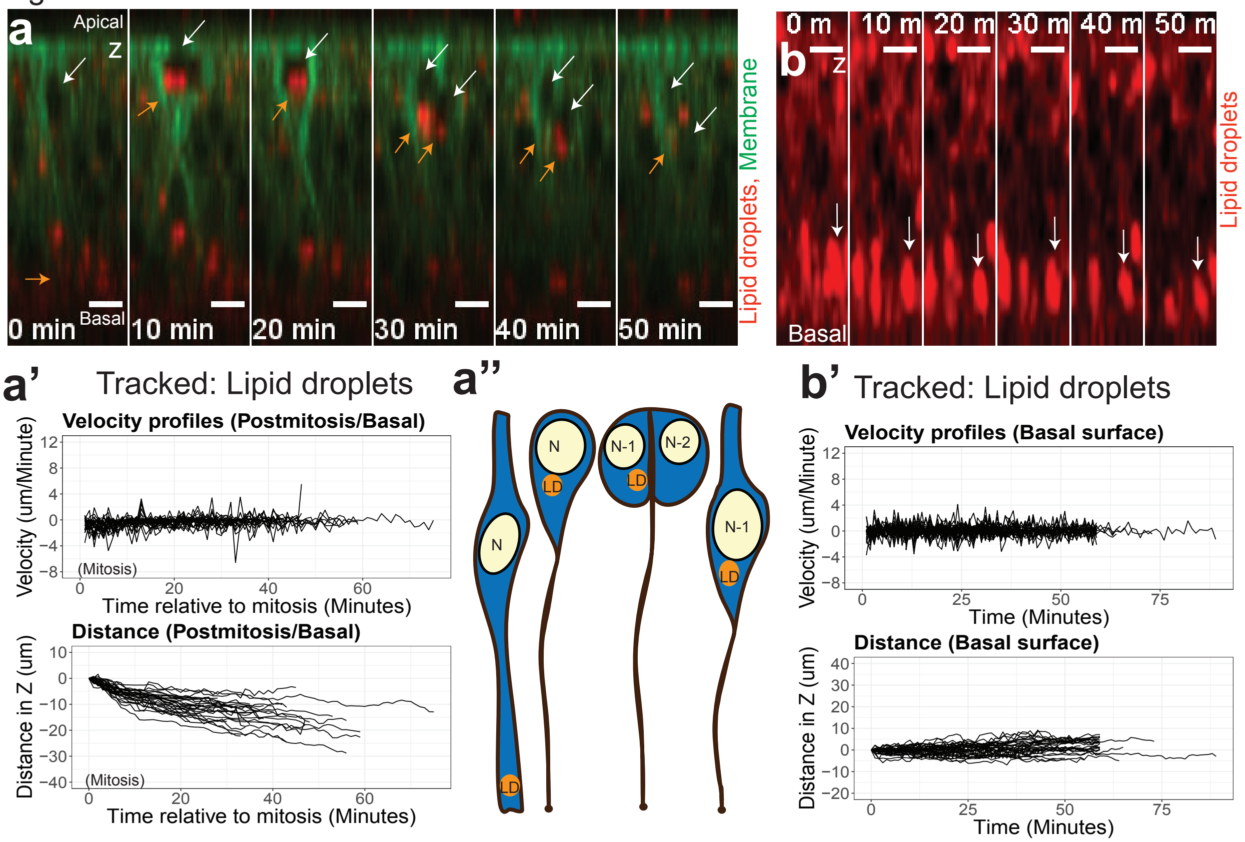


Fig S3: An apical to basal flow following mitosis and ground state of lipid droplet motion at basal surface. (a, b) *c765*-GAL4>UAS-*PLCdelta-PH-GFP* with 1:2000 LipidTOX Deep Red. (a) Representative images for rapid flow of lipid droplets (orange arrow) before mitosis and a much slower apical to basal flow afterwards with the nucleus. Nucleus (white arrow). (b) Ground state of the dynamics of LDC at basal surface. (a', b') Quantification of velocity and distance profiles of tracked LDC in the apical-basal axis. (a') Basal-directed flows (n=30 LDC). (b') Stochastic motion of lipid droplets (n=37 LDC). (a'') Model of type I flow, mitosis and basal-directed flow. Scale bars: 5 µm.

Fig S4

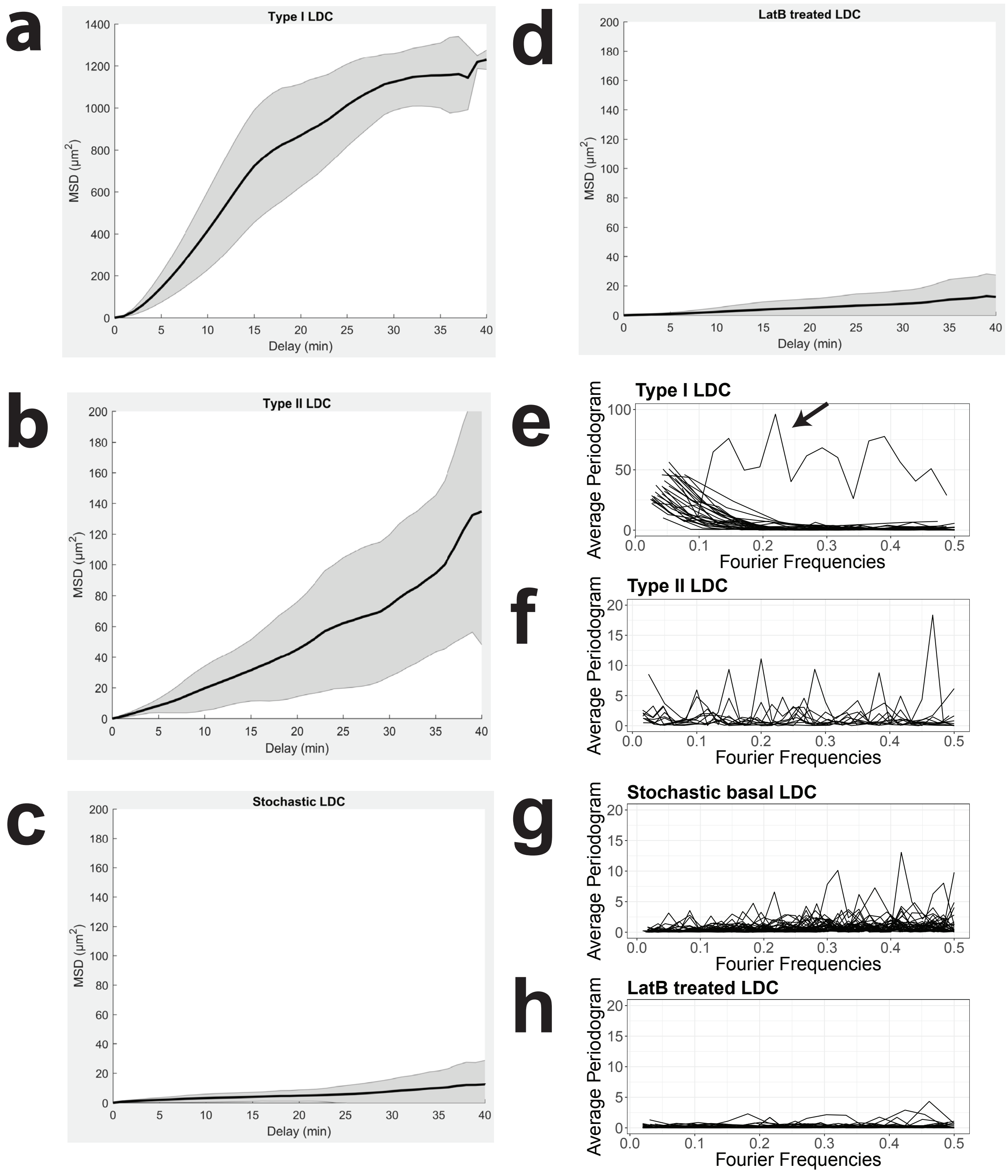
a
Type I LDC
MSD (µm2)
Delay (min)
b
Type II LDC
MSD (µm2)
Delay (min)
c
Stochastic LDC
MSD (µm2)
Delay (min)
d
LatB treated LDC
MSD (µm2)
Delay (min)
e
Type I LDC
Average Periodogram
Fourier Frequencies
f
Type II LDC
Average Periodogram
Fourier Frequencies
g
Stochastic basal LDC
Average Periodogram
Fourier Frequencies
h
LatB treated LDC
Average Periodogram
Fourier Frequencies

Fig S4: MSD analysis and frequency analysis (a) LDC type I flow is advection. (b) Type II LDC motion is advection. (c) LDC ground state at basal surface in non-mitotic cells is stochastic. (d) LatB treated LDC motion is stochastic. (e) Average periodogram for LDC type I flow with a large low frequency components. Arrow represents an outlier sample which had relevant period of type I motion and underwent mitosis, the tracking data revealed large periods of resting motion during which there is unusually large movements which may be due to tracking error. (f) Average periodogram for LDC type II flow. (g) Average periodogram for stochastic LDC. (h) Average periodogram for LatB treated LDC.

Fig S5

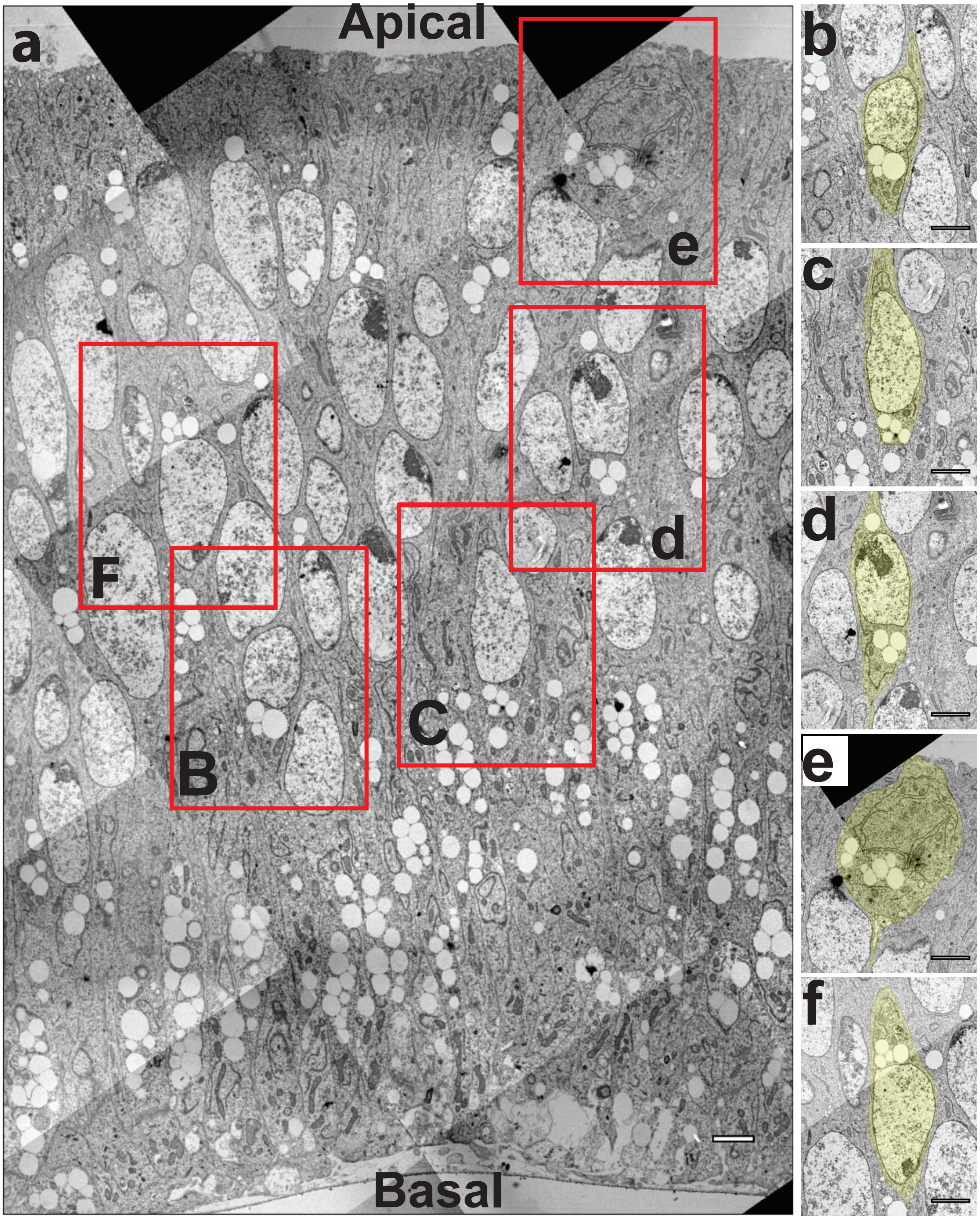

Fig S5: Nuclear morphologies and nucleus proximity to lipid droplets as an indicator of the flow. (a) TEM stitched section of columnar epithelium cells with pointed droplet nuclear morphology. (b-d) Pointed droplet morphology of the nucleus is towards the apical surface representing apical-directed flow. (b-c) Represents nucleus in type II flow from the tightly packed lipid droplets cluster and nucleus is much closer to basal surface. (d) Represents a type I flow from the tightly packed lipid droplet cluster since nucleus is closer to the mitotic zone. (b, d) Nucleus is deformed at on the side of the lipid droplets representing the increased hydrostatic pressure under the nucleus. (e) Rounded mitotic cell in the apical mitotic zone with organelles inside. (f) Represents a cell in basal-directed flow from the pointed droplet morphology towards basal surface and the lipid droplets on the apical side of the nucleus. Scale bars: 5 µm.

Fig S6

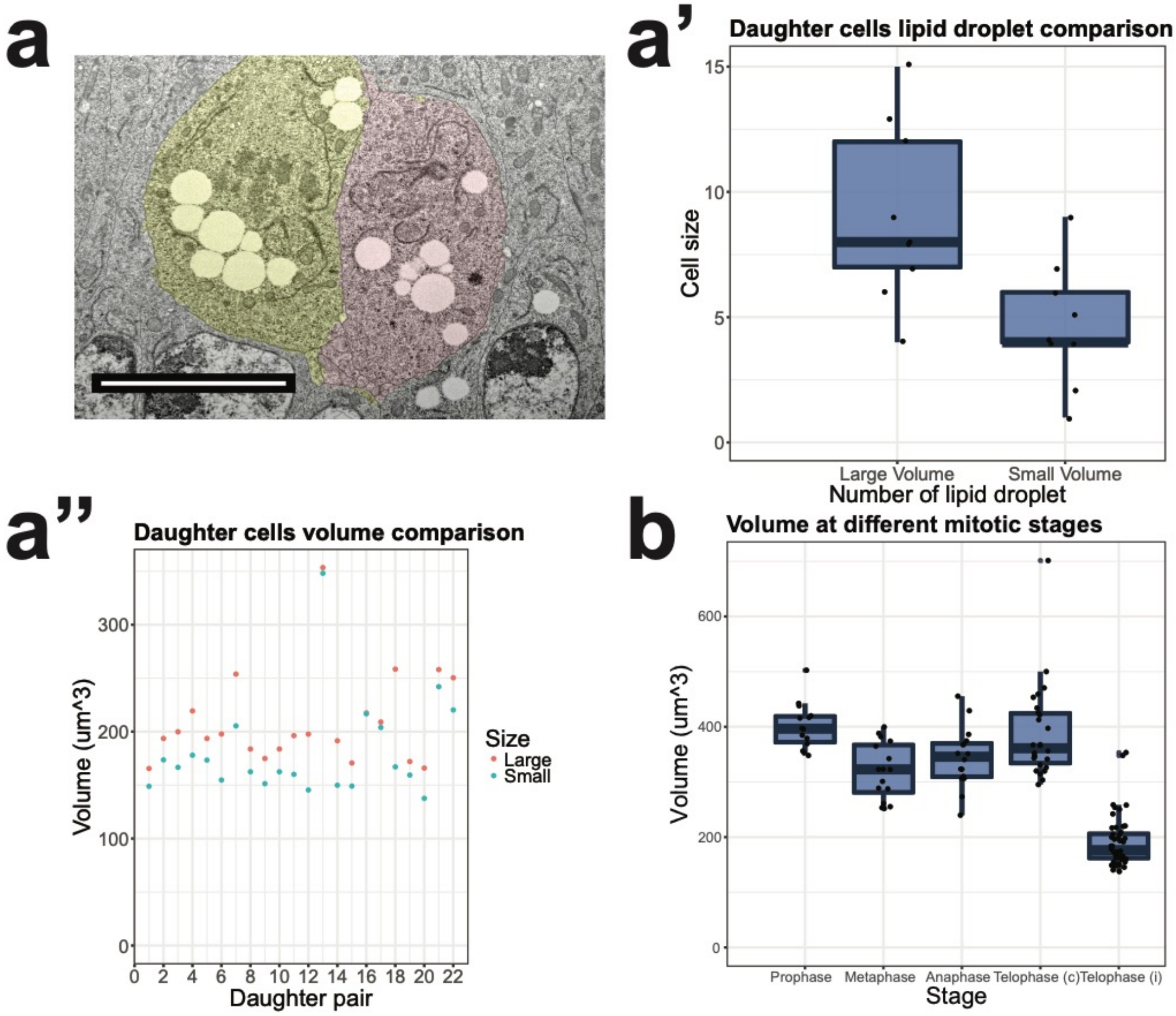


Fig S6: Lipid droplet/volume distribution during telophase, volume of the cells and 3D nucleus and lipid droplet sizes. (a) Represents TEM section of the late telophase nucleus to illustrate distribution of lipid droplets (Yellow, pink are the two daughter cells. (a') Lipid droplet distribution among daughters counted in 3D from confocal images. (a') Volume distribution of among daughter pairs, calculated in 3D from confocal images. (b) Volume of cells at different mitotic stages, Telophase (c) is combined volume of the cell(s) and Telophase (i) is volumes of individual cells. Scale bar: 5 µm.

**Supplementary Movies**

Supplementary Movie S1: Time-lapse confocal microscopy movie of lipid droplets and mitochondria in 3D in mitotic cells.

Supplementary Movie S2: Time-lapse confocal microscopy movie illustrating the model of the type I apical-directed flow. **Scale bars**: 5 µm.

Supplementary Movie S3: Time-lapse confocal microscopy movie illustrating the type II flow type I flow followed by post-mitosis flow, type I mitochondrial flow and resting motion of lipid droplets. **Scale bars**: 5 µm.

Supplementary Movie S4: Time-lapse confocal microscopy movie illustrating the drug treatment in xy slice and z slice. **Scale bars**: 5 µm.
